# Impact of pubertal onset on region-specific *Esr2* expression

**DOI:** 10.1101/2021.03.31.437916

**Authors:** Carly M. Drzewiecki, Elli P. Sellinger, Janice M. Juraska

## Abstract

In female rats, pubertal onset is associated with maturation of the medial prefrontal cortex (mPFC) and mPFC-mediated behaviors. These behavioral and anatomical changes are likely due to effects of estrogen at the nuclear estrogen receptor beta (ERβ), which is expressed at higher levels than the estrogen receptor alpha (ERα) isoform in the adult mPFC. Researchers have previously quantified ERβ protein and *Esr2* RNA in rodents during early postnatal development and adulthood, but an adolescent-specific trajectory of this receptor in the mPFC has not been documented. Given that levels of *Esr2* can fluctuate in the presence or absence of estrogens, puberty and the subsequent rise in gonadal hormones could influence ERβ expression in the adolescent brain. To further explore this, we used RNAscope to quantify the amount of *Esr2* mRNA in pre-pubertal adolescent, recently post-pubertal adolescent, and adult female rats. We show here that *Esr2* expression decreases significantly in the mPFC, striatum and motor cortex between pre-pubertal adolescence and adulthood. In the mPFC, this decrease occurs rapidly at pubertal onset, with no significant decrease in *Esr2* levels between the recently post-pubertal and adult cohort. In contrast in the striatum and motor cortex, there were no significant differences in the amount of *Esr2* between pre- and post-pubertal females. Insofar as the amount of *Esr2* is proportional to functional ERβ, these results suggest ERβ decreases in a region-specific pattern in response to pubertal onset and highlight a role for this receptor in the maturational events that occur in the female rat mPFC at puberty.

## Introduction

Adolescence, the transitional period between childhood and adulthood, is often defined by pubertal onset and the subsequent rise in gonadal hormones. These hormones play an important role in the development and reorganization of the adolescent brain^1^, particularly within the female medial prefrontal cortex (mPFC)^2^. For instance, ovarian hormones are necessary for the neuronal pruning that occurs in this region during adolescence^3^. Pubertal onset also coincides with synaptic pruning as well as an abrupt decrease in the number and density of perineuronal nets (PNNs) ^4,5^, and pubertal estrogens drive an increase in inhibition in the frontal cortex^6^. Together, these results suggest pubertal onset could rapidly influence cortical plasticity in the female mPFC. A possible functional correlate of these changes is an induction at pubertal onset of adult-like behaviors on a cognitive flexibility paradigm ^7^, a task that relies on GABAergic functioning in the prefrontal cortex ^8^. Given that all these developmental events occur at pubertal onset when females are experiencing a rapid increase in estrogen, estrogen receptors most likely play an important regulatory role in these processes.

Estrogen binds with equal affinity to its two known nuclear receptors: the estrogen receptor α (ERα) and the estrogen receptor β (ERβ) ^9,10^. Both receptors are present throughout the human and rodent brain, as well as peripheral tissue, though their expression patterns are often region- and tissue specific. For example in adult rats, ERα mRNA is more highly expressed in the pituitary, uterus and other peripheral reproductive tissues ^11^, and both receptors are detected in the hypothalamus in nuclei-specific patterns ^12^. As such, ERα has often been implicated in reproductive behaviors and fertility ^13,14^. ERβ is more prevalent in the rodent cortex and hippocampus ^12,15–17^, and plays an important modulatory role in cognitive ^18,19^, social ^20^, and emotional behaviors ^21–25^ These effects are likely due in part to ERβ’s influences on synaptic plasticity ^23,26–28^.

Most of the research on ERβ levels has been conducted in adult, ovariectomized subjects, though studies have demonstrated that the amount of ERβ RNA fluctuates during development, particularly within the mPFC. At birth and across the first postnatal week, ERα is the prominent receptor subtype in the rodent mPFC ^29,30^, but this pattern soon reverses, and ERβ becomes the dominant subtype in the juvenile mPFC ^12,30,31^. The same is true in the adult cortex, with ERα being virtually absent in the adult mPFC, but work that examines how levels of ERβ fluctuate during adolescence and pubertal onset specifically is lacking. Both brain and peripheral ERβ expression levels are altered in the presence or absence of both endogenous and exogenous estrogens in region- and dose-specific patterns ^11,32–35^. Therefore, the rapid increase in endogenous estrogens at pubertal onset could influence ERβ expression in the mPFC, a region we have previously shown is sensitive to these hormones during this time^2^

Currently, there is no validated antibody that binds to ERβ using immunohistochemistry in perfused tissues ^36^. The newer technological method RNAscope, however, has been employed to visualize *Esr2*, ERβ mRNA, in the rodent brain ^37^. Here we aimed to quantify levels of *Esr2* in female adolescent and adult rats, with particular emphasis on the effects of pubertal onset on the amount *Esr2*. We measured *Esr2* levels in the mPFC, as well as the dorsal striatum and motor cortex, two regions that have low levels of Esr2 and ERβ in adulthood ^12,17,38^. Levels of *Esr2* were quantified before and after pubertal onset in adolescent females and in early adulthood.

## Methods

### Subjects

Subjects were the female offspring of Long Evans rats ordered from Envigo [Formerly Harlan] (Indianapolis, IN) that were bred in the vivarium of the Psychology Department at the University of Illinois. Rats were weaned at P25, housed in groups of two or three with same-sex littermates and provided food and water *ad libitum* with a 12:12 light-dark cycle. All procedures were approved and adhered to guidelines set forth by the University of Illinois Institutional Care and Use Committee.

Two of the timepoints for sacrifice in this study were determined based on pubertal status, so pubertal onset was assessed daily beginning at postnatal day (P) 29 in females. Pubertal onset was determined by vaginal opening, which corresponds to increases in luteinizing hormone and estrogen ^39^. Within a litter, when subjects reached puberty, they were sacrificed 24 hours later with an age- and sex-matched sibling that had not yet reached puberty. These are referred to as the “Post” and “Pre” groups, respectively. The average age at sacrifice for the Pre and Post groups was 34.55 days with a range of 32–37. A third group of litter-matched females was sacrificed at P60, referred to here as the “Adult” group.

At sacrifice, subjects were deeply anesthetized with a lethal injection of sodium pentobarbitol and perfused intracardially with a 0.1 M solution of phosphate buffered saline (PBS) followed by a 4% paraformaldehyde fixative solution in PBS (pH 7.4). Brains were post-fixed in the paraformaldehyde solution for an additional 24 hours followed by immersion in a cryoprotectant 30% sucrose solution. At this point all brains were coded to conceal group from the experimenters. After 72 hours, brains were sliced in 40 µm coronal sections using a freezing microtome. Sections were stored at -20°C in an anti-freeze storage solution until processing (30% glycerol, 30% ethylene glycol, 30% dH_2_0, 10% 0.2 M PBS). Sections of this tissue were also used in Drzewiecki et al.^5^.

### Fluorescent in Situ Hybridization with RNAscope

Levels of *Esr2*, the gene that encodes ERβ, were assessed using RNAscope fluorescent *in situ* hybridization (FISH). The *Esr2* probe (ACD, catalog #317161) was verified *in silico* by ACD and recently by Kanaya et al.^37^. Additionally, the probe was further verified with the positive probe (PPIB, provided by ACD), and absence of signal with the negative probe (provided by ACD) as well as in slices where the *Esr2* probe was excluded. RNAscope has been proven to be an effective method in perfused brain slices that are 40 µm thick^40^.

The RNAscope protocol for fixed tissue from Advanced Cell Diagnostics (ACD) was followed for visualization or the *Esr2* probe, with minor modifications as advised by the company. The protocol was conducted over two days. To begin day 1, tissue slices stored in anti-freeze storage solution were washed triplicate in PBS and mounted on Superfrost Plus slides and allowed to dry completely at room temperature. The slides were then baked at 60° C for 30 minutes, followed by incubation in ice cold 4% paraformaldehyde (made the day prior and at a pH 7.4) for 15 minutes at 4° C. Slides were further dehydrated using a series of ethanol dilutions, and then baked again for 30 minutes at 60° C. Next, slides were incubated in hydrogen peroxide for 10 minutes at room temperature before beginning the target retrieval process.

The target retrieval solution provided by ACD was diluted to 1X and warmed to at least 99° C along with dH_2_O in a steamer. The slides were placed in the warm water solution for 10 seconds and then quickly moved to the warmed target retrieval solution where they were incubated in the steamer for 5 minutes. After retrieval, slides were briefly rinsed in dH_2_O at room temperature, followed by a 5 minute incubation in 100% ethanol. Finally, slides were air dried at room temperature and a hydrophobic barrier was drawn around slices using ImmEdge Hydrophobic Barrier Pen (ACD, catalog # 310018).

Following target retrieval, all steps were performed using chemicals provided in the Multiplex Fluorescent Reagent Kit v2 (ACD, catalog # 323100), and all incubations were done in a humidity controlled chamber at 40° C. First, slides were coated in the protease solution and incubated for 30 minutes. Then slides were rinsed twice for two minutes in dH_2_O and incubated in the *Esr2* probe (ACD, catalog #317161) for 2 hours, followed by a wash step. After hybridization of the probes, all wash steps were conducted using wash buffer solution (ACD, catalog # 310091), and consisted of two washes in the buffer for two minutes each. The slides were stored overnight in a 5X saline-sodium citrate (SSC) buffer solution.

The following day, slides underwent a wash step and began incubations in the three separate amplification solutions. Each amplification was separated by a wash step. After the amplification steps, channel 1 was established and the slices were incubated in a 1:750 solution of Opal 570 (Akoya Biosciences, catalog # FP1488001KT) diluted in TSA Buffer (ACD). Following incubation in the fluorophores, slices were washed and then incubated in HRP channel blocker. Finally, slides were briefly rinsed with DAPI, before coverslipping with ProLong Gold Antifade Mountant (ThermoFisher, catalog # P36930). Slides were dried in the dark overnight at room temperature, and then stored at 4° C until imaging.

### Microscopy, Imaging, and Fluorescence Analysis

Slides were imaged using a Zeiss LSM 880 microscope within 3 days of the FISH protocol. Images were acquired from layer II/III of the mPFC, layers II/III and V/VI of the motor cortex, and the dorsolateral and dorsomedial striatum. Regions of interest were identified using the DAPI stain, and 8 separate images (4 per hemisphere) were acquired for each subject within each subregion analyzed (Figure 1). Images were obtained by taking z-stacks of 30 separate images across 20.3 µm of tissue at 8 unique sites within each region. The z-stacking specifications were held consistent across all subjects.

**Figure 1.**
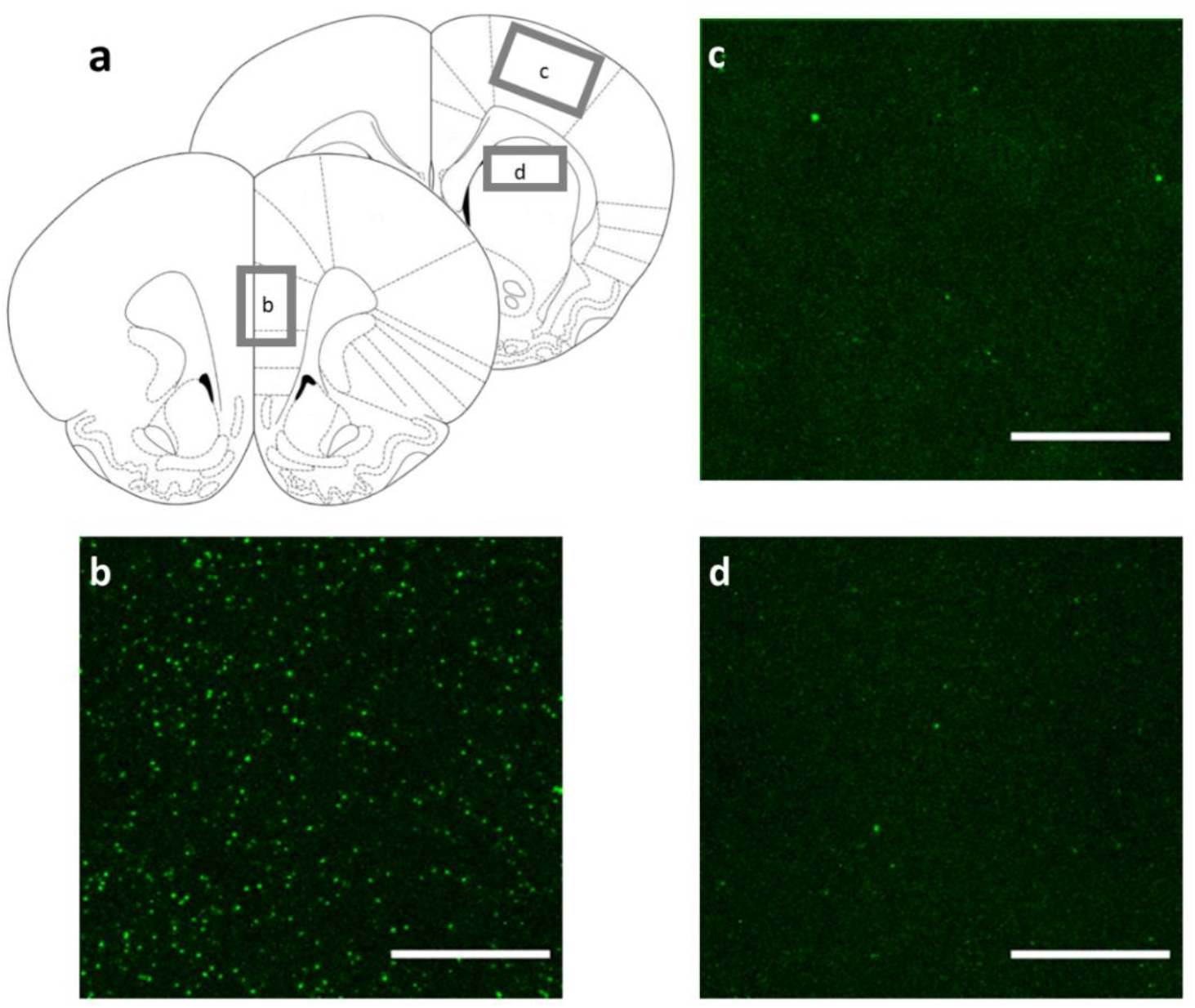
*Sampling regions (a) and representative images of* Esr2 *puncta in the mPFC (b), motor cortex (c), and dorsal striatum (d). Images show* Esr2 *puncta in pre-pubertal, adolescent females. Scalebar = 20 µm*.

An image processing macro was written for ImageJ (NIH), which allowed the images to be consistently and quickly analyzed. The macro first projected the z-stacks into one image. Then each image was converted to grayscale, and the rolling ball function was used to subtract background noise. Finally, a threshold of 75 arbitrary units (aus) was applied, and all pixels that met this threshold were quantified and recorded. Each image represented 369,693.5 µm^3^ of tissue, and the total number of puncta per image was averaged within each subject.

### Statistical Analysis

All statistical tests were conducted using RStudio. Because each region had to be imaged at a unique laser power setting due to region-specific background staining levels, all three regions sampled were analyzed separately. Mixed linear effects models were performed using the “lmerTest” package for the average number of puncta within each image with pubertal group as a fixed factor and litter included as a random factor. Then, separate tests for analysis of variance (ANOVA) were conducted on the models and the residuals were checked for normality using the Shapiro-Wilk test. When residuals from the ANOVA models were not normally distributed, the non-parametric Kruskal-Wallis test was used for analysis. Post hoc testing for parametric data was accomplished using the “emmeans” package with a Bonferroni correction for 3 comparisons, and post hoc testing for the non-parametric data from the motor cortex and striatum was conducted with the Dunn test for multiple comparisons with a Bonferroni correction for 3 comparisons.

Data from 9 pre-pubertal females, 12 post-pubertal females, and 8 adults was included in this study, and 1,152 images were analyzed from the 29 total subjects. mPFC images from one pre-pubertal female had to be excluded due to an imaging error.

## Results

In the mPFC, there was a significant effect of group on puncta density (*F* = 9.60, *p* = 0.001). Post hoc testing demonstrated significant differences between the pre- and post-pubertal females (*p* = 0.002) and the pre-pubertal and adult subjects (*p* = 0.01) (Figures 2a & 3).

**Figure 2.**
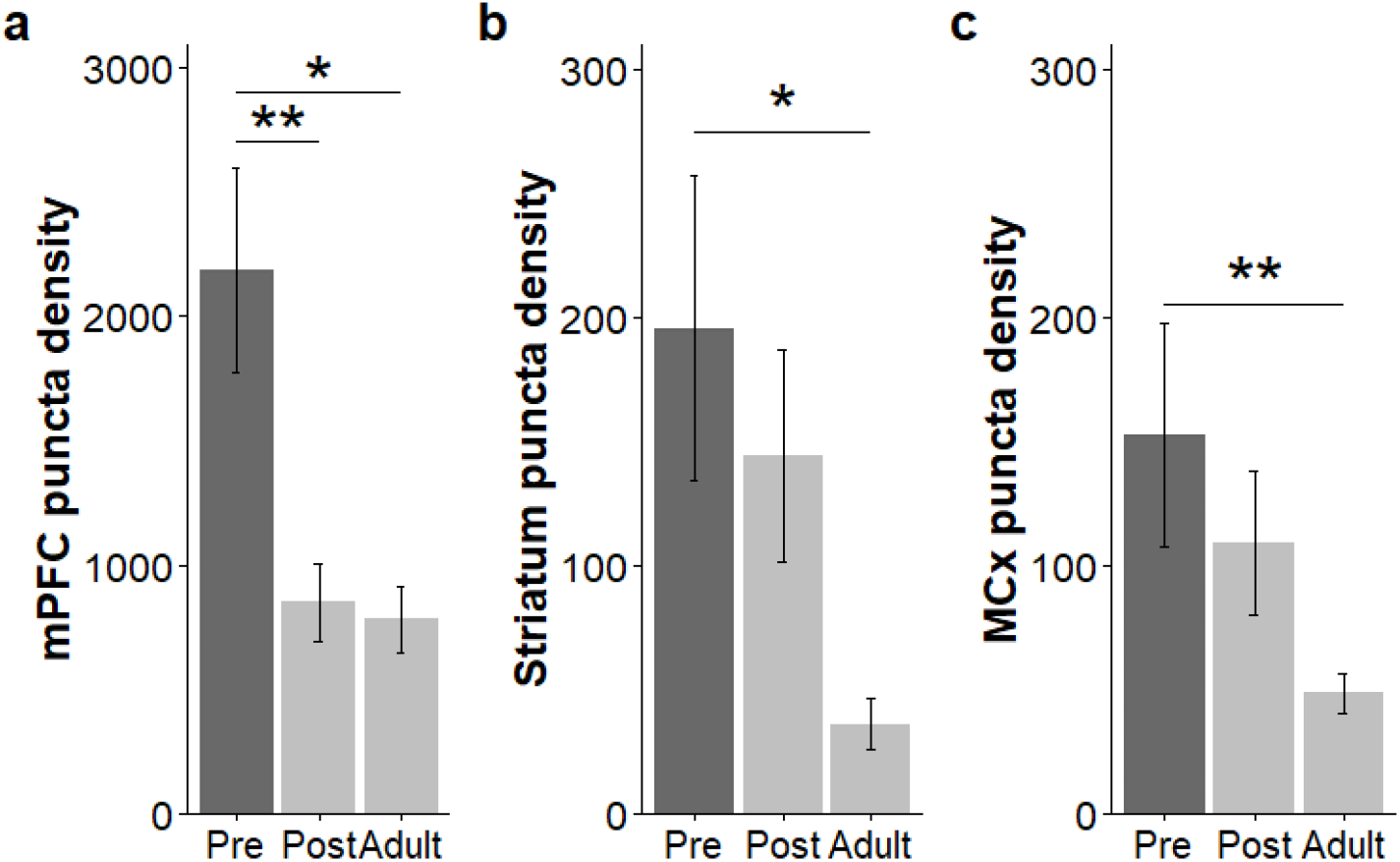
*The density of* Esr2 *puncta was significantly decreased in the mPFC (a) after pubertal onset and in adulthood when compared to the pre-pubertal group. In both the striatum (b) and the motor cortex (MCx) (c), the puncta density was significantly lower in adult subjects compared to pre-pubertal subjects, with no significant decreases following pubertal onset. In all plots, lighter bars demarcate subjects that had reached puberty at the time of sacrifice. Darker bars indicate pre-pubertal subjects. [* p < 0.05; ** p < 0.01]*

**Figure 3.**
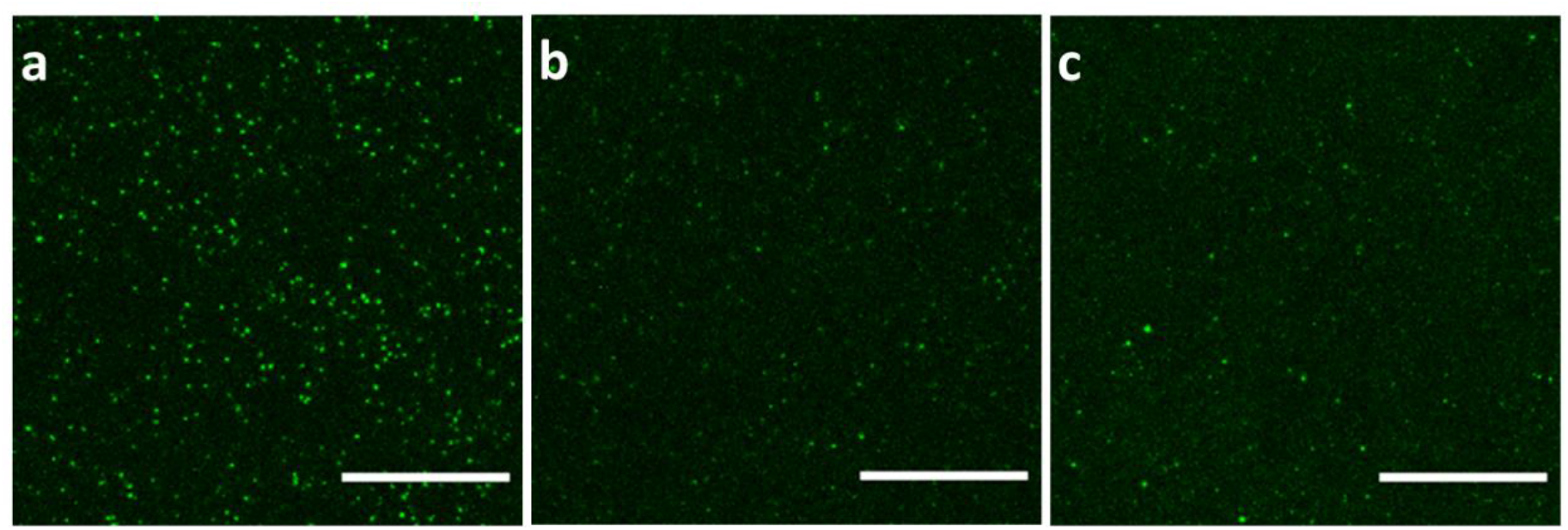
*Representative images from the mPFC.* Esr2 *puncta density was significantly higher in pre-pubertal females (a) compared to recently post-pubertal (b) and adult (c) subjects. Puncta density between post-pubertal adolescents and adults did not significantly differ. Scalebar = 20 µm.*

There was no significant effect of subregion in the striatum on the average number of puncta per image (data not shown), so the data was collapsed across both areas for subsequent analysis. There was a significant effect of group on the total number of puncta (*χ*^*2*^ = 7.46, *p* = 0.02). A post hoc revealed a significant difference between the pre-pubertal and adult females (*p* = 0.03) (Figure 2b).

Likewise, within the motor cortex, no differences in the density of puncta per image were found between the upper and lower cortical layers (data not shown), so all layers were combined for analysis. There was a significant effect of group on puncta density (*χ*^*2*^ = 9.80, *p* = 0.007), and the post hoc test again revealed a significant difference between the pre-pubertal and adult females (*p* = 0.002) (Figure 2c).

## Discussion

*Esr2* was quantified in pre- and post-pubertal, as well as adult females. In the three neural regions examined, *Esr2* decreased between pre-pubertal adolescence and adulthood. However, in the mPFC, the amount of *Esr2* dropped to adult-like levels just 24 hours after pubertal onset; this abrupt decrease between pre- and post-pubertal subjects was not observed within the dorsal striatum or the motor cortex. We have previously hypothesized that gonadal hormones act to reorganize the mPFC during adolescence, and these findings support this in that changes to *Esr2* may roughly translate to changes in levels of ERβ. Also, by using age-matched pre- and post-pubertal female siblings, we demonstrate that pubertal status, rather than linear age, can be a better predictor of cortical *Esr2* expression in adolescent females.

Many studies have shown that both endogenous and supplemental estrogen can influence *Esr2* levels in various cortical and subcortical regions, though the direction of these effects are highly region- and most likely dose-specific. For example, within the rodent hypothalamus, both ERα and ERβ vary in concentration across the estrus cycle ^41–43^. Estrogen supplementation increases *Esr2* in the arcuate nucleus but decreases *Esr2* mRNA expression in the paraventricular nucleus, amygdala, entorhinal cortex and thalamus ^33,34,44^. Some studies show that estrogen supplementation decreases *Esr2* in the bed nucleus of the stria terminalis ^45,46^, while others show no changes ^34^, indicating that the effects of estrogen on *Esr2* levels are not always consistent. Nonetheless, ERβ and *Esr2* expression are clearly dynamic and can fluctuate in the presence or absence of estrogen.

Interestingly, several studies have shown that estrogen does not alter *Esr2* levels in the adult mPFC^44,46^, suggesting that our observed decrease in *Esr2* at pubertal onset was not driven by the presence of estrogen. However, it is important to note these studies examined adult females that had been ovariectomized after pubertal onset and supplemented with a consistent dose of estrogen meant to replicate levels seen during proestrus. Given that the effects of estrogen on behavior and estrogen receptor expression can be highly dose-specific ^33,47^, studies that artificially replace estrogen after surgical removal of the ovaries likely do not mimic natural conditions.

Additionally, the aforementioned studies that examine the effects of estrogen on *Esr2* were all conducted in adult subjects, and recent work suggests the functional effects of estrogen on the prefrontal cortex may be age-specific. For example, estrogen enhances fear extinction via actions in the infralimbic cortex in adult rats^19,48–50^, though the opposite is true in recently post-pubertal adolescent females, and pubertal estrogens impair fear extinction in these subjects ^51^. Additionally, recently post-pubertal females show decreased mPFC PNNs compared to age- and litter-matched pre-pubertal females. This decrease persists for at least a week of adolescence before increasing to adult-like levels at P60 ^5^, suggesting that circulating estrogens in post-pubertal females act differently on PNNs than circulating estrogens in adulthood. Therefore, it is possible that circulating estrogens decrease *Esr2* in the developing mPFC, while having no effect on *Esr2* levels in the adult mPFC.

Studies that exclude adolescence and only examine juvenile and adult subjects have suggested that *Esr2* levels remain consistent across the lifespan ^15,52^, but our results demonstrate *Esr2* decreases between adolescence and adulthood in all three regions examined. In the prefrontal cortex, ERβ mRNA begins to rise shortly after birth and continues to increase until at least P25 ^29,30^. Our study documents an abrupt decrease in prefrontal *Esr2* at puberty, suggesting that ERβ may follow an inverted-U shaped trajectory and reach peak levels in the adolescent mPFC just before pubertal onset. If true, this would mirror our finding that the total number of synapses in the female mPFC peaks around pubertal onset ^4^, and further demonstrate the plastic nature of the adolescent cortex ^2^.

It is less clear if the same developmental pattern of *Esr2* occurs in the adolescent striatum or motor cortex. Despite the fact that both regions are directly modulated by estrogens ^24,53,54^, little evidence exists about how the hormonal environment influences ERβ expression or ontogeny in the motor cortex or striatum. We did not observe an abrupt decrease in *Esr2* in either region at pubertal onset, which may be due, in part, to the fact that there were fewer *Esr2* puncta in these regions. This is consistent with what others have reported^12,38^and lower amounts of *Esr2* could make it difficult to detect subtle differences in *Esr2* expression. However, it is worth noting that there was a significant decrease in *Esr2* between pre-pubertal adolescence and adulthood, so our results do not entirely exclude a role for post-pubertal estrogens in *Esr2* expression in these regions. Future research on the topic is warranted.

Though we have previously reported female rats lose a significant number of neurons in the mPFC around the time of pubertal onset ^55^, we do not hypothesize that the abrupt decrease in prefrontal *Esr2* shown in this study is a direct result of this neuronal pruning. We describe here a nearly 50% decrease in *Esr2* puncta between pre- and post-pubertal females, which is significantly larger than the 11% decrease in neurons after puberty reported by Willing & Juraska ^55^. Additionally, within the mPFC, ERβ co-localizes extensively with inhibitory parvalbumin (PV) cells ^56^. Though the density and intensity of PV cells does decrease in the female mPFC between P40 and P70^57^, the magnitude of this drop is again far lower than the 50% decrease observed here in *Esr2* puncta. Our observed decrease in *Esr2* at pubertal onset appears to be due to a downregulation of the receptor and not simply driven by cellular pruning.

Given the anatomical relationship between ERβ and GABAergic cells ^16,56^, a decrease in prefrontal *Esr2* at pubertal onset could have implications for cortical inhibition. Fine tuning of the excitatory/inhibitory balance in the developing cortex is crucial for maturation of inhibitory control during adolescence ^58^, and pubertal estrogens organize mature inhibitory transmission in the mouse frontal cortex ^6^. Additionally, cortical inhibition in naturally-cycling adult female rats is regulated by the action of estrogen at ERβ ^59^. Our results add to a growing body of literature that suggests pubertal onset in females mediates an important shift in inhibitory neurotransmission.

Using RNAscope, we have found *Esr2* levels are rapidly downregulated in the mPFC at pubertal onset. We demonstrate the decrease in *Esr2* between adolescence and adulthood is not unique to the mPFC and was also observed in the dorsal striatum and motor cortex. While previous studies have quantified ERβ mRNA in juvenile or ovariectomized adult animals, we add a unique focus on puberty and adolescence to previous findings and demonstrate that this receptor is very much in flux across the pubertal transition. We hypothesize that levels of *Esr2* may peak pre-pubertally, but future work that includes juvenile and older adult cohorts is necessary to fully elucidate the ontogeny of this receptor. Nonetheless, the results presented here underscore the importance of accounting for pubertal onset when studying adolescence and further highlight a critical role for estrogen in maturation of the female mPFC.

## Acknowledgements

This work was funded by National Institute of Environmental Health Sciences Grant P01 ES002848-Project3 (to JMJ). EPS was partially supported by National Institute of Environmental Health Sciences Grant T32 ES007326. We thank the Core Facilities at the Carl R. Woese Institute for Genomic Biology for use of the LSM 880 microscope, as well as Austin Cyphersmith for his technical guidance.

